# Different hydrological conditions after permafrost thaw result in distinct microbial community compositions

**DOI:** 10.1101/2024.12.28.630578

**Authors:** Muhammad Waqas, Patrick Liebmann, Cordula Vogel, Milan Varsadiya, Haitao Wang, Olga Shibistova, Milan Novák, Tim Urich, Georg Guggenberger, Jiří Bárta

## Abstract

Permafrost degradation leads to the formation of contrasting hydrological conditions such as water-saturated anoxic and dry oxic soils, which significantly influence the bacterial community structure and abundance. To investigate the bacterial abundance and diversity under these hydrological conditions, we collected soil samples from different horizons of both dry and wet degraded permafrost soils and non-degraded intact permafrost soil for comparison. The bacterial alpha diversity, measured by the Observed and Chao1 indices, was significantly greater in wet degraded permafrost soil (wet site) and intact permafrost soil (intact site) than in dry degraded permafrost soils (dry site). Notably, the wet and intact sites exhibited similar levels of alpha diversity as well as shared greater number of zOTUs. The relative proportion of most bacterial taxa was significantly differed among the sites. At the class level, dry site was dominated mainly by K-strategic bacteria like aliphatic degraders (Thermoleophilia), acidophilic and cellulolytic (Acidobacteriae), while wet site was dominated by mainly r-strategic bacteria from class Gammaproteobacteria and anoxygenic aerobic phototrophs (Gemmatimonadetes). According to Spearman correlation analysis, the relative proportion of bacterial taxa in dry and wet sites showed significant correlation with soil physicochemical parameters, whereas fewer correlations were found in intact site. The relative proportion of Pseudomonadota classes and Acidobacteriota were positively correlated with SOC, N, and C:N ratio, but were negatively correlated with pH in both dry and wet sites. In contrast, anaerobic methylotrophs (Methylomirabilota), anoxygenic photoheterotrops (Chloroflexota), Gemmatimonadota and filamentous Actinobacteriota were negatively correlated with SOC, N, and C:N. Additionally, strong correlations were observed between bacterial taxa and extracellular enzyme activities, where Alphaproteobacteria, Gammaproteobacteria, and Acidobacteriota showed positive correlations with both hydrolytic and oxidative enzymes in dry and wet sites, except for PerOx in wet site where they showed negative correlation. Conversely, Gemmatimonadota and Actinobacteriota displayed negative correlations with enzymes in dry and wet site, except for the PerOx in wet site, where they showed positive correlation.

## Introduction

Permafrost soil represent an important microbial ecosystem (Waldrop et al., 2023) and a globally significant reservoir of sequestered carbon (C) (Schuur et al., 2022). Despite being regarded as a harsh environment, permafrost regions host diverse microbial communities that exhibit remarkable resilience and adaptability to extreme environment (Abramov et al., 2021; Walker et al., 2006). Upon thawing of permafrost, these long-preserved microbial communities can quickly resume or enhance their metabolic functions, influencing nutrient availability and contributing to shifts in ecosystem processes (Feng et al., 2020a; Y. Li et al., 2024; Romanowicz et al., 2023). Consequently, there is growing concern regarding the microbial mineralization of sequestered organic carbon (OC) in frozen ground in response to climate change (Anisimov & Zimov, 2021; DeConto et al., 2012; Koven et al., 2011; Schuur et al., 2008; Yokohata et al., 2020).

Above the perennially frozen ground, shallow seasonally thawed active layer hosts diverse microbial communities and are more abundant compared to those present in the below frozen layers (Chen et al., 2017). These microbial communities in active layer are metabolically active (Jansson & Taş, 2014; Wilhelm et al., 2011), responding to seasonal changes in temperature and moisture (Schostag et al., 2019), and driving key biogeochemical cycles during the thawed months (Pautler et al., 2010). Descriptions of active layer microbial communities have been reported from diverse environments and show a high level of variation in diversity and abundance. Among the diverse microorganisms found, bacteria are the most dominant group with dominant phyla including a wide range of copiotrophs, oligotrophs, phototrophs and acidophiles depending on the permafrost soil conditions and play significant role in biogeochemical processes. For example, in Alaskan permafrost soils, copiotrophic decomposers of Pseudomonadota, and actinobacteria played significant roles in soil organic matter (SOM) turnover, particularly in nutrient-rich and moist tundra environments (Mackelprang et al., 2011). In Western Canadian permafrost soils, dominant groups include SOM decomposers (Pseudomonadota), actinobacteria (Actinobacteriota), acidophiles (Acidobacteriota), and anoxygenic phototrophs (Chloroflexota), crucial for N cycling and SOM degradation (Varsadiya et al., 2021), while in Siberian permafrost, acidophiles (Acidobacteriota) are abundant due to adaptation to the acidic, nutrient-poor conditions (Kobabe et al., 2004). Bacterial community structure is shaped by multiple environmental factors, including the rapid freeze-thaw cycles, moisture content, pH, temperature fluctuations, oxygen levels, and SOM quality (Ganzert et al., 2014; Schostag et al., 2019; Siciliano et al., 2014; Tytgat et al., 2016). With rising global temperatures, these environmental factors increasingly influence bacterial activity and community composition, particularly in relation to SOM decomposition and the emission of CO_2_ and CH_4_ (Jansson & Taş, 2014; Jin & Ma, 2021; Mackelprang et al., 2011; Schuur et al., 2015).

Rising global temperatures have become a critical concern, especially in permafrost regions, where an estimated 2°C increase could reduce Northern Hemisphere permafrost areas by over 50% within the next century (Karjalainen et al., 2020) . The early signs of this transformation are already evident, as permafrost ground temperatures have increased by approximately 1°C since the 2000s (Bartsch et al., 2023). This steady warming, driven by increasing global temperatures is contributing to increase in the thickness of active layer and permafrost degradation across vast regions (Jin & Ma, 2021; G. Li et al., 2022; Smith et al., 2022). These changes can influence the bacterial communities by shifting the environmental factors like changing in aboveground vegetation, enhancement of plant rooting depth, water content, and release of significant pool of sequestered OC (Feng et al., 2020b; Liu et al., 2022; M.-H. Wu et al., 2022). However, there remains limited understanding of how bacterial communities transform when permafrost thawing leads to distinct hydrological conditions like dry aerated or wet anoxic degraded permafrost soils.

In degraded permafrost environments with well-drained conditions due to better drainage and evapotranspiration, the landscape often supports deeper rooting vegetation systems, along with higher nutrient levels and increased oxygen availability (Jia et al., 2020; Jorgenson et al., 2001; Ren et al., 2018a). These dry and aerated conditions promote greater bacterial diversity and abundance, as well as increased enzymatic activities, as the high respiration rates have been previously reported in these conditions in response to warming, which is linked to the decomposition of SOM (Natali et al., 2015). On the other hand, increased bacterial activity under dry and aerated conditions, can also enhance soil development through the production of extracellular polymeric substances (EPS), which play a crucial role in the stabilization and aggregation of soil particles (Costa et al., 2018). These EPS not only contribute to the physical binding of soil particles, leading to the formation of aggregates, but may also help foster OM within these aggregates, thereby protecting from decomposition (Opfergelt, 2020; Totsche et al., 2018). This contrasts with wet and anoxic conditions where oxygen is limited, and can significantly alter bacterial communities, hampering the growth of aerobic bacteria that thrive in oxygen-rich environments (Jia et al., 2020). In such environments, the OC can be more stable in the short term due to reduced bacterial activity. However, this condition can still a favourable environment for anaerobic bacteria and facilitate anaerobic processes such as denitrification and methanogenesis (Varsadiya et al., 2021; Wagner et al., 2009). Overall, these contrasting hydrological conditions of permafrost degradation landscapes can drive significant shifts in bacterial community composition, favouring aerobic bacteria in dry and oxic soils and anaerobic in wet anoxic, ultimately shaping bacterial processes and their impact on OC dynamics. We hypothesize that (i) the enhanced aeration and deeper root systems present in well-drained dry permafrost degradation landscape leads to a richer and more abundant of bacterial community, outperforming the bacterial resilience observed in water-saturated wet landscape that mimic intact permafrost; (ii) that bacterial taxonomic richness in water-saturated degraded permafrost soils will be less diverse, aligning closely with that of intact permafrost, indicating that excessive moisture may limit microbial adaptability and diversity as compared to their drier counterparts, and (iii) the interplay between soil physicochemical factors and bacterial relative abundance will reveal greater correlations in degraded permafrost soils of both dry and wet suggesting that environmental conditions drive microbial composition more substantially than in intact permafrost ecosystems. For this, we investigated two types of hydrologically degraded permafrost soils landscapes alongside non-degraded intact permafrost soils for total bacterial community and their relationship with soil physicochemical parameters. All study sites were located within one kilometre near the Fairbanks, Alaska, USA.

## Materials and methods

### Study sites and sampling

The research area is located in the Interior Alaska, USA, near the city of Fairbanks. The detailed description of the study sites and sampling is reported in the Liebmann et al. (2024). In brief, three landscapes types including one intact permafrost soil (intact site) and two degraded permafrost soil sites (dry and wet sites) were studied to examine the effects of permafrost thaw on bacterial structure in relation to soil physicochemical properties. The annual ground temperature ranges from -2.5°C to -0.5°C for the intact site, and from 1.9°C to 2.7°C for the dry and wet sites. At each site, four soil pits were prepared with a depth of 100 cm (dry and wet sites) or until reaching the permafrost level in 45-55 cm (intact site) for sampling the defined organic layer, mineral topsoil and subsoil within the active layer. The topsoil depth increment was consistently defined as 2-7 cm below the organic layer for all three sites. In the degraded permafrost soils (dry and wet sites), the subsoil increment was set at 45-50 cm below the organic layer. For the intact site, where this subsoil depth lies within the frozen ground, the subsoil sampling was adjusted to a 5 cm increment above the permafrost table, approximately 10-15 cm below the topsoil depth, staying within the active layer. The soil materials were divided into three sample sets, which were used for soil physicochemical analysis, including EPS extraction and enzyme activity analysis. For the genetic material extraction, approximately 3 g of soil sample were immersed with 2 volumes of LifeGuard Soil Preservation Solution (Qiagen, Hilden, Germany) immediately after sampling. All Lifeguard-treated samples were constantly kept at 4°C until processing. In this study, a total of four replicates (one from each pit) of both topsoil and subsoil were used for each study site, while organic layer samples included three replicates from the intact site, one from dry site, and two from wet site.

### Soil physicochemical parameters

Moisture levels were determined by drying the reweighed samples at 60 °C for 48 hours, followed by reweighing. The determination of SOC, Nitrogen (N), pH and Base saturation (BS) was carried out as outline by Liebmann et al. (2024) using established procedures.

### EPS extraction

EPS from the soil samples was extracted using the cation exchange resin (CER) method described in Redmile-Gordon et al. (2014). The procedure began with the preparation of phosphate-buffered saline (PBS) and CaCl_2_ solutions, which were pre-cooled to 4°C. CER was washed using twice its weight in PBS and then filtered through glass wool to remove excess liquid. In skirtless centrifuge tubes, 2.5 g of soil was added, ensuring the tubes were securely capped to minimize evaporation. For the extraction, 25 mL of pre-cooled 0.01M CaCl_2_ was added to the soil samples, which were then incubated on shaker at 4°C for 30 minutes to extract easily soluble OM. Following centrifugation at 3200 x g for 30 minutes, the supernatant was carefully decanted, and the pellets were retained for EPS extraction. The pellets were mixed with CER and PBS in the same tubes and incubated horizontally on a shaker at 4°C for 2 hours. After incubation, the tubes were centrifuged at 4200 x g for 20 minutes to separate the soil and resin from the supernatant. The resulting supernatant, now containing the EPS were then lyophilised (Freeze dryer, Christ Beta 1-8 LSC plus) for 48 hours to obtain the dry weight of EPS. The content of EPS-polysaccharides was determined using the phenol-sulfuric acid method as described by DuBois et al. (1956), with glucose serving as the standard. The EPS-protein concentration was determined using the modified Lowry method, employing bovine serum albumin as the standard (Frølund et al., 1996). The EPS-sugar as measured as glucose equivalent and EPS-protein were then calculated per gram of soil.

### Enzymatic activities

Hydrolytic and ligninolytic enzymes involved in geochemical processes were determined using the standard method described in Bárta et al. (2010, 2014). One gram of soil was suspended in 50 mL of distilled deionized nuclease-free water (ddH2O) and ultrasonicated at low energy (120 W) for 4 minutes. For β-glucosidase, cellobiosidase, chitinase, leucine aminopeptidase, and phosphatase, fluorometric techniques with 4-methylumbelliferyl (MUF)-linked substrates for the glycosidic and phosphatase enzymes, and 7-amino-4-methylcoumarin (AMC) as a substrate for leucine aminopeptidase (50–300 µM) were employed. A 200 µL sample of the soil suspension were pipetted into black microtiter plates in 3 analytical replicates. Then 50 µL of the MUF or AMC labeled substrates were added. For each sample, a standard curve with MUF or AMC was measured for the calibration. Plates were incubated in the dark for 30 minutes, and the first fluorescence was measured at 465 nm emission with 360 nm excitation (Tecan Infinite F200 fluorimeter, Männedorf, Switzerland). Fluorescence was measured again after 60 and 120 minutes. Enzyme activities were expressed in nmol g⁻¹ dry weight of soil h⁻¹. To assess phenoloxidase (PhOx) and peroxidase (PerOx) activities (ligninolytic enzymes), microtubes were prepared in triplicates by adding 1 ml of soil suspension to 1 ml of 5 mM L-3,4-dihydroxyphenylalanine (L-DOPA) substrate dissolved in 50 mM acetate buffer. Negative controls for the sample contained 1 ml of soil suspension mixed with 1 ml of acetate buffer, while negative controls for the substrate contained 1 ml of acetate buffer combined with 1 ml of the substrate. For PerOx assays, an additional 100 μl of 0.3% H_2_O_2_ was added to each tube. The microtubes were covered with aluminium foil and incubated on an orbital shaker for 1 hour at 20°C. Enzyme activity was quantified spectrophotometrically at 460 nm. Both PhOx and PerOx activities were measured separately and expressed in μmol h⁻¹ g⁻¹ dry weight of soil.

### DNA extraction and Barcode amplicon pair end sequencing of prokaryotic 16SrRNA genes

For the total genomic DNA extraction, the Lifeguard-treated soil samples were used according to the recommended protocol. The LifeGuard solution was removed via centrifugation and total DNA was extracted using the PowerSoil DNA elution kit (Qiagen, Hilden, Germany). The final elution volume was 50 μL and the eluted DNA was stored at -20°C. The total genomic DNA extracts from the soil samples were sent to SEQme sequencing company (Prague, Czech Republic) for the preparation of a library and sequencing using the MiSeq2500 platform. Library was prepared using the Earth Microbiome Project (EMP) protocol with modified universal primers 515FB/806RB (Caporaso et al., 2011). Bacterial 16S rRNA raw pair-end reads (250 bp) were joined and quality filtered using USEARCH v.11 to obtain reads approx. 250 bp in length (Edgar, 2013). Amplicons were then trimmed to equal lengths. Bacterial unique reads were grouped into zero-radius OTUs (zOTUs) using a UNOISE 3.0 algorithm which also included chimera check and removal (Edgar & Flyvbjerg, 2015). Taxonomic assignment of each bacterial zOTUs was performed using the BLAST algorithm (min E-value=0.001, min similarity 99%) against the curated ARB Silva 138.2 database (Quast et al., 2012). All unassigned zOTUs and those assigned to Chloroplast and Mitochondria were discarded. Raw fastq 16S rRNA gene sequences of the whole microbiome were uploaded to European nucleotide archive (ENA) under the study n. PRJEB79237.

### Statistical Analyses

For the data analysis, a phyloseq object was created by importing the zOTUs table, taxonomy, and metadata, using the “Phyloseq” package in R. The number of shared and unique zOTUs among the dry, wet and intact sites and different horizons was determined using the “VennDiagram” package. The alpha diversity was calculated using observed number of species, Chao1, and Shannon using phyloseq package. One-way analysis of variance (ANOVA) followed by Tukey’s honestly significant difference (HSD) test or Wilcoxon rank-sum test were used to determine the differences in alpha diversity indices, the relative abundance of major bacterial phyla/class, and the soil physiochemical parameters. The *P*-values were adjusted by Bonferroni. The bacterial community composition pattern in dry, wet and intact sites and different horizons were analyzed by nonmetric multidimensional scaling (NMDS) using the Vegan package. Analysis of similarities (ADONIS) and permutation multivariate analysis of variance (PERMANOVA) were performed to test the statistically significant differences among bacterial communities with Bray-Curtis distances and 999 permutations. The normality of bacterial relative abundance and chemical parameters were checked using Shapiro-Wilk test before the comparative analysis. The relative abundance was standardized by Hellinger data transformation using decostand function. Soil chemicals parameters were standardized with z-score transformation. Redundancy analysis (RDA) with permutation test for constrained correspondence analysis was used to identify the effects of soil properties on the bacterial community. The forward selection was used to identify the most significant soil parameters to explain the pattern of bacterial communities. Spearman’s correlation analysis was used to estimate the relationship between the bacterial relative abundance at phyla/class level and soil physiochemical properties. All the plots were generated using the ggplot v 3.4.4 package.

## Results

### Soil physiochemical properties

Soil physicochemical properties differed between the studied sites and different horizons of each site. The level of moisture and pH, along with the abundance of SOC, N, EPS-protein and the C:N ratio was higher in the intact site compared to the dry and wet sites. Though these differences across the sites were not significant (Table S1). Moisture and the amount of SOC, N, C:N ratio, EPS-sugar and EPS-protein were decreased from the organic layer to the mineral topsoil and further to the mineral subsoil, across the all three sites (Significant differences were only between topsoil and subsoil, Table 1). Similarly, BS also decreased with increasing depth in the wet and intact sites, but in the dry site it was increased from the topsoil to subsoil. In contrast to the other parameters, pH increased from the organic layer to the topsoil and further to the subsoil in all three sites. Compared to the horizons across the sites, the amount of SOC and N was greater in the mineral topsoil of the dry and wet sites than the intact site. While the SOC and N was higher in the subsoil of the intact site than the dry and wet sites. Similar patterns were also followed for the EPS-glucose and EPS-protein (Table 1). They were higher in the topsoil of the dry and wet sites and lower in the subsoil compared to the intact site.

**Table 1:** Soil physicochemical properties per horizon of dry, wet and intact sites.

| Landscape | Horizon | Count | Moisture<br>[%] | pH | BS<br>[%] | SOC<br>[mg g<sup>-1</sup>] | N<br>[mg g<sup>-1</sup>] | C:N | EPS_sugar<br>[μg g<sup>-1</sup>] | EPS_protein<br>[μg g<sup>-1</sup>] |
| --- | --- | --- | --- | --- | --- | --- | --- | --- | --- | --- |
| Dry | Organic Layer | 1 | 81.4 | 4.2 | 61.2 | 400.7 | 15.8 | 25.3 | na | na |
|  | Topsoil | 4 | 32.2±7.1 (a) | 4.5±0.2 (a) | 26.6±5.7 (a) | 28.9±7.3 (a) | 1.5±0.4 (a) | 19.7±1.3 (a) | 80±23.6 (a) | 45.1±15.1 (a) |
|  | Subsoil | 4 | 22.1±4.7 (a) | 6.3±0.1 (b) | 66.5±1.3 (b) | 3.7±0.8 (b) | 0.4±0.1 (b) | 9±1 (b) | 36.7±16.5 (b) | 15.2±3 (b) |
| Intact | Organic Layer | 3 | 88.8±5.6 (a) | 5.5±0.9 (a) | 89±3.3 (a) | 374.6±28.1 (a) | 11.7±2.6 (a) | 33.5±9.6 (a) | Na | Na |
|  | Topsoil | 4 | 44.2±20.8 (b) | 5.9±0.1 (a) | 45.7±2.6 (b) | 23.9±5.6 (b) | 1.6±0.4 (b) | 15.4±0.5 (b) | 73.4±27.9 (a) | 45.7±39.7 (a) |
|  | Subsoil | 4 | 33.2±7 (b) | 6±0.1 (a) | 44.6±5.4 (b) | 16.6±8.4 (b) | 1.2±0.6 (b) | 13.7±0.7 (b) | 64.6±35.3 (a) | 23.2±11.3 (a) |
| Wet | Organic Layer | 2 | 75.4±5.6 | 4.4±0.9 | 72±4.6 | 347.2±93.5 | 11.4±0.9 | 30.8±10.6 | na | na |
|  | Topsoil | 4 | 40.6±11.6 (a) | 5.8±0.7 (a) | 65.2±9.4 (a) | 37±16 (a) | 1.9±0.7 (a) | 19.2±2.2 (a) | 210.6±150.8 (a) | 53.8±31.8 (a) |
|  | Subsoil | 4 | 30.1±4.7 (a) | 6.1±0.2 (a) | 51.8±1.4 (b) | 4.3±1.8 (b) | 0.5±0.1 (b) | 9±1.7 (b) | 58.9±17.4 (a) | 18.6±11.1 (a) |

Averages and standard deviation were shown. The significant difference between horizons was calculated by One-Way ANOVA and followed by Tukey's HSD test.
Different letters in the brackets indicate a significant difference. Count: number of replicates; na, no data available. BS: Base saturation, SOC: Soil organic carbon, N:
Nitrogen. SOC and N data were obtained from Liebmann et al. (2024) , and were integrated into the current analysis to correlate with bacterial community composition.

The activities of extracellular enzymes specifically β-glucosidase, cellobiosidase, chitinase, leucine aminopeptidase, phosphatase, and PhOx exhibited substantial variability across dry, wet, and intact sites (Table S2). Notably, enzyme activities were highest in the dry site, followed by the wet site, while the lowest activities were recorded in the intact site, although the differences between the intact and other sites were not statistically significant. Contrary, PerOx had significantly the greatest activity in the intact site then the dry and wet sites (Table S2). Enzymatic activities varied significantly across soil horizons at each site, with data highlighting differences primarily between the topsoil and subsoil (Table 2). All investigated enzymes exhibited greater activity in the organic layer of the dry and wet sites, with the exception of PerOx, which showed lower activity in the organic layer compared to the mineral horizons. When comparing topsoil and subsoil within each site, β-glucosidase, chitinase, and phosphatase activities were consistently higher in the topsoil across all three sites. Furthermore, when comparing horizons across the sites, the activities of β-glucosidase, cellobiosidase, chitinase, phosphatase, and PhOx were greater in the topsoil of the dry and wet sites than in the intact permafrost soil. In contrast, these enzymes exhibited higher activity in the subsoil of the intact site compared to the dry and wet sites.

**Table 2:** Enzyme abundance per horizon of dry, wet and intact landscape.

| Locality | Horizon | Count | β-glucosidase<br>[nmol<br>g<sup>-1</sup> dw h<sup>-1</sup>] | Cellobiosidase<br>[nmol<br>g<sup>-1</sup> dw h<sup>-1</sup>] | Chitinase<br>[nmol<br>g<sup>-1</sup> dw h<sup>-1</sup>] | Phosphatase<br>[nmol<br>g<sup>-1</sup> dw h<sup>-1</sup>] | Leucinaminopeptidase<br>[nmol<br>g<sup>-1</sup> dw h<sup>-1</sup>] | PhOx<br>[μmol h<sup>-1</sup><br>g<sup>-1</sup>dw] | PerOx<br>[μmol h<sup>-1</sup> g<sup>-1</sup>dw] |
| --- | --- | --- | --- | --- | --- | --- | --- | --- | --- |
| Dry | Organic Layer | 1 | 1802.1 | 656.1 | 1335.3 | 3494.2 | 34 | 625.3 | 688.9 |
|  | Topsoil | 4 | 170.5±117.2 (a) | 27.5±20.8 (a) | 54.3±21.9 (a) | 181±64.3 (a) | 15.6±11.8 (a) | 114.6±34.2 (a) | 790.7±34.4 (a) |
|  | Subsoil | 4 | 10.9±3 (b) | 6.7±1.3 (a) | 6.6±2.7 (b) | 30.1±5.3 (b) | 15.5±3.9 (a) | 72.5±27.6 (a) | 692.4±84.3 (a) |
| Intact | Organic Layer |  | na | na | na | na | na | na | na |
|  | Topsoil | 4 | 81.3±15.1 (a) | 14.5±3.9 (a) | 35.4±7.7 (a) | 90.4±17.4 (a) | 22.7±6.2 (a) | 101.5±32.9 (a) | 1099.9±191 (a) |
|  | Subsoil | 4 | 36.1±9.9 (b) | 10.8±3.4 (a) | 17±5.4 (b) | 51.2±19.7 (a) | 19.3±6.9 (a) | 73±5.8 (a) | 1066.4±114.6 (a) |
| Wet | Organic Layer | 2 | 686.4±623.4 | 60.4±47 | 193.3±24.6 | 988.6±313.6 | 63.8±46.8 | 194.1±83.7 | 600.1±599.7 |
|  | Topsoil | 4 | 150.4±78.4 (a) | 14.1±8.3 (a) | 57.3±29.9 (a) | 189.7±100.9 (a) | 89.2±47.8 (a) | 179.5±88.9 (a) | 1113.5±296.3 (a) |
|  | Subsoil | 4 | 9.2±3.8 (b) | 5±2.4 (a) | 6.3±2.3 (b) | 21.2±4.3 (b) | 3.4±2.1 (b) | 93.9±22.2 (a) | 1265.8±153.8 (a) |

Averages and standard deviation were shown. The significant difference between horizons was calculated by One-Way ANOVA and followed by Tukey's HSD test.
Different letters in the brackets indicate a significant difference. Count: number of replicates: na: no data available.

### Bacterial diversity and richness

The taxonomic assignment of pre-processed high-quality sequences yielded 3930 bacterial zOTUs and 15 archaeal zOTUs at the 99% sequence similarity cut-off. The number of unique bacterial zOTUs observed in the dry, wet and intact sites varied from 181 to 278 (Figure 1). The number of unique zOTUs found in the dry site was 278, while it was 176 and 181 zOTUs in the wet and intact sites respectively. About 1789 (46%) zOTUs were found common to all the sites and considered as “core microbiome”. A higher proportion of zOTUs (26%) was common between the wet and intact sites while only 3.4% common zOTUs were shared between the dry and intact sites. The similarities and differences of bacterial zOTUs were further explored between the horizon of each studied site. The proportion of common bacterial zOTUs was found higher (39%) between the horizons of the intact site, followed by the horizons of the wet site (25%) (Figure 1). The least shared zOTUs were found between the horizons of the dry site (9.7%). The number of unique zOTUs was higher in the organic layer of wet (378) and intact (355) sites compared to the dry sites, where the number of the unique zOTUs was found higher in the subsoil (787). The highest proportion of the common zOTUs was among the topsoil and subsoil of wet (29%) and intact (27%) sites, while in the dry site the proportion of common zOTUs was between the organic layer and topsoil (24%) (Figure 1). The similarities and differences in bacterial zOTUs were also examined across comparable horizons of the studied sites (Figure S1). The proportion of common zOTUs was higher between the horizons of wet and intact sites. Topsoil of the wet site shared 30% zOTUs with the topsoil soil of intact site and subsoil of wet site shared 28% of zOTUs with the subsoil of intact site. The least shared zOTUs was among the topsoil of the dry site with the topsoil of the intact site (6.4%) and subsoil soil of dry site with subsoil soil of intact site (5.3%).

**Figure 1:**
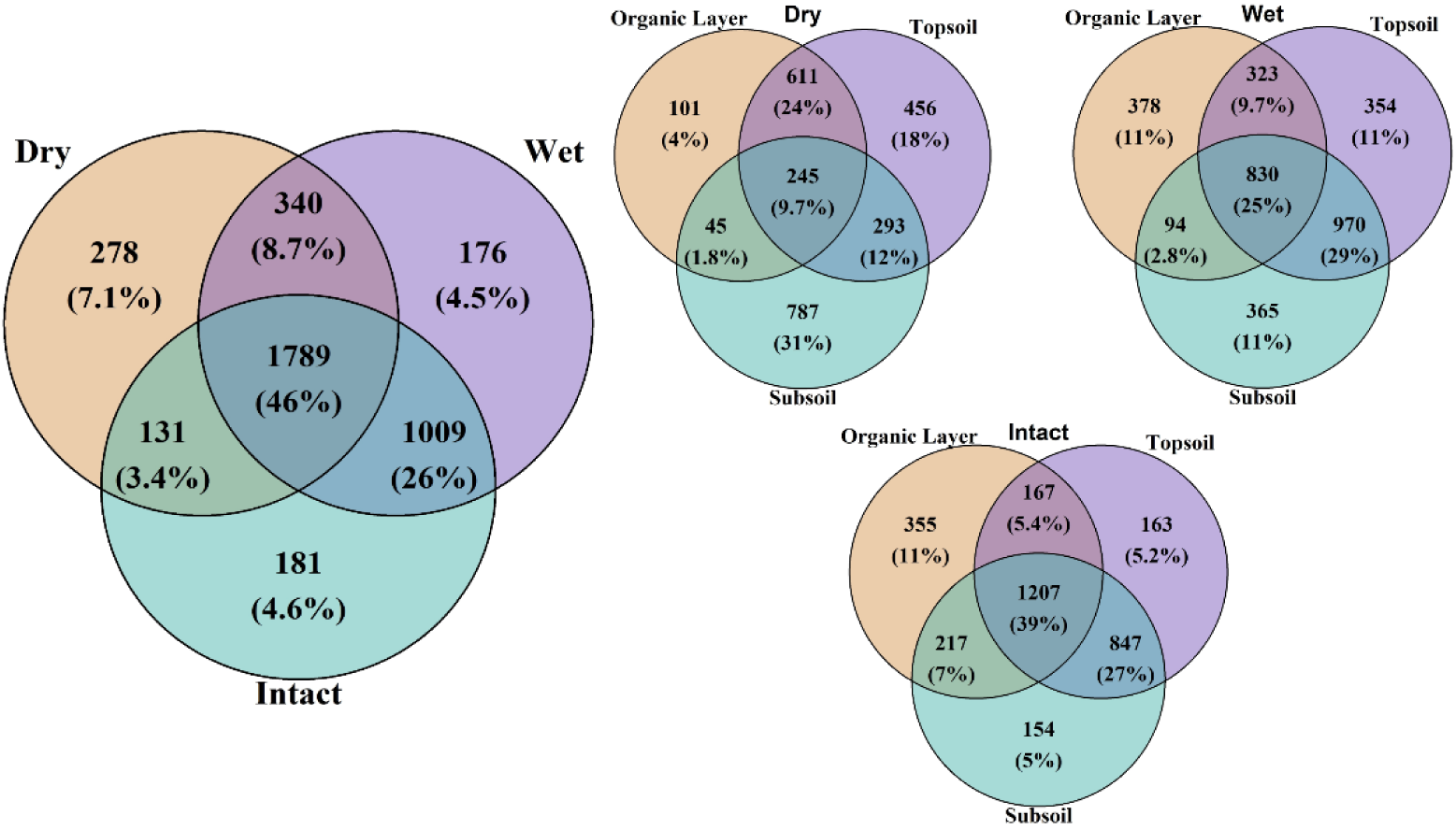
Venn diagram illustrating the overlap of zOTUs across the dry, wet, and intact sites and per horizons of each site. The diagram displays both the raw counts and the percentages of zOTUs unique to each condition and shared among them, with the sizes of the circles representing the total number of zOTUs in each site/horizon. The intersections highlight the shared zOTUs between the sites/horizons of each site, with the central area representing the zOTUs common to all three site/among the horizon of each site, i.e. core microbiome.

The diversity, richness and evenness of the bacterial community was statistically determined based on the observed number of species, Chao1, and Shannon indices. Species richness and Chao1 index was highest for the intact and wet sites compared to the dry site (Figure 2). The Shannon evenness was higher for the intact site while the dry and wet sites showed the least. The significant differences were observed for the diversity richness and Chao1 index between the dry and intact sites, as well as between dry and wet sites. No significant difference was observed among intact and wet sites (Figure 2). Alpha diversity, assessed by observed species and Chao1, was highest in topsoil of dry site, where as in the wet and intact sites it was highest in the subsoil, although all three horizons did not show significant differences (Table 3).

**Figure 2:**
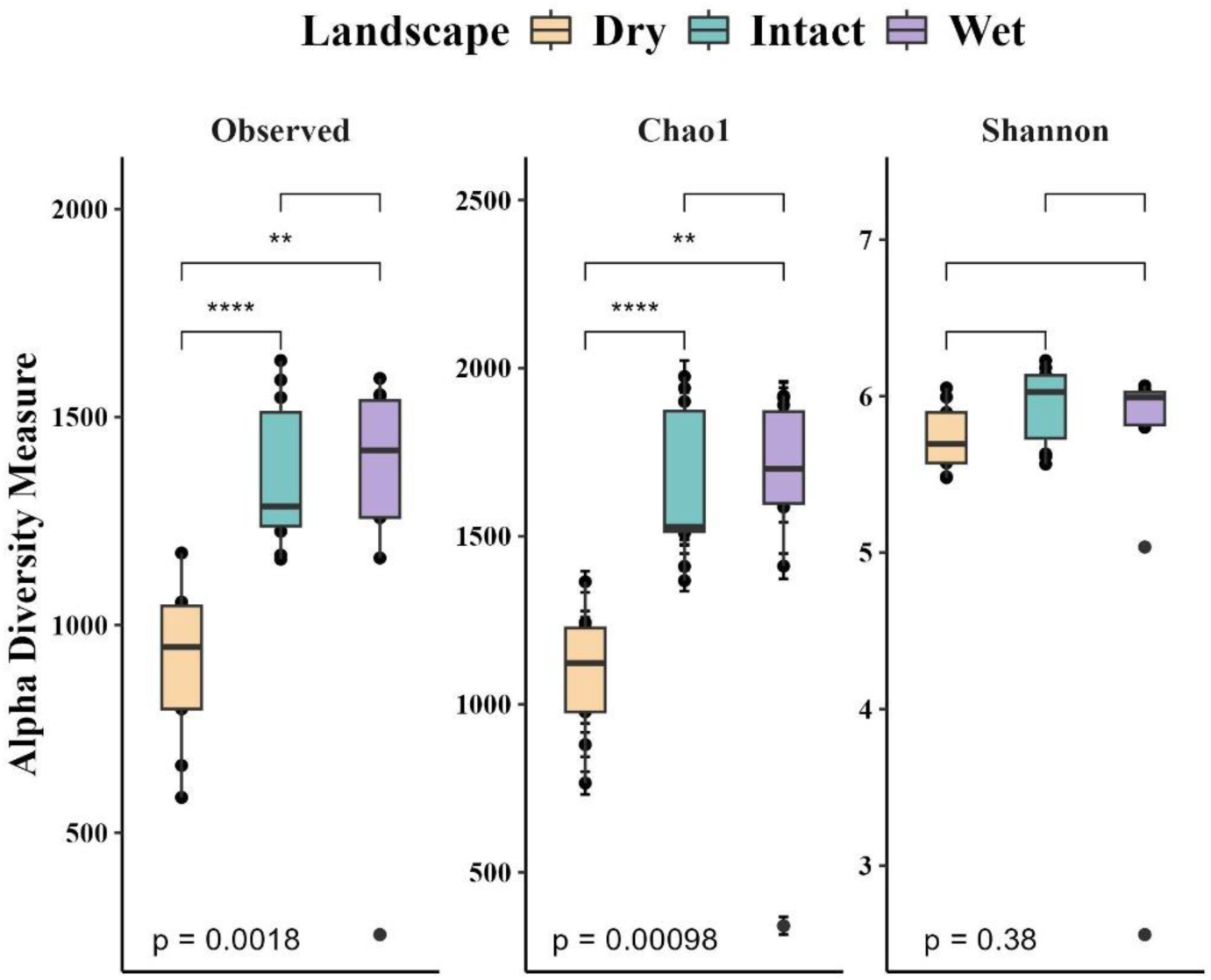
Observed species richness (alpha diversity) of the total bacterial community across dry, wet, and intact sites. The boxes represent the interquartile range (IQR) with the median shown as a line within the box. Whiskers extend to 1.5 times the IQR, and outliers are shown as individual points. An overall One-Way ANOVA was performed to assess the differences in diversity across the sites, with the resulting *p-value* displayed at the bottom of the plot. Statistical comparisons between the landscapes were performed using the Wilcoxon rank-sum test with Benjamini-Hochberg correction for multiple testing. Significant differences between the sites are indicated as follows: * p < 0.05, ** p< 0.01, *** p < 0.001, and **** p < 0.0001. Non-significant comparisons are not shown.

**Table 3:**
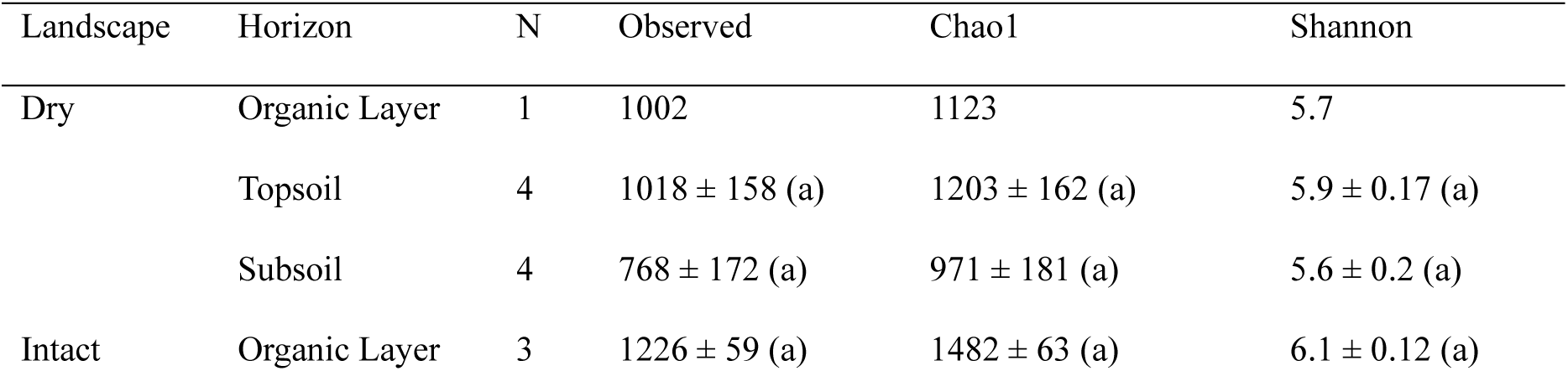
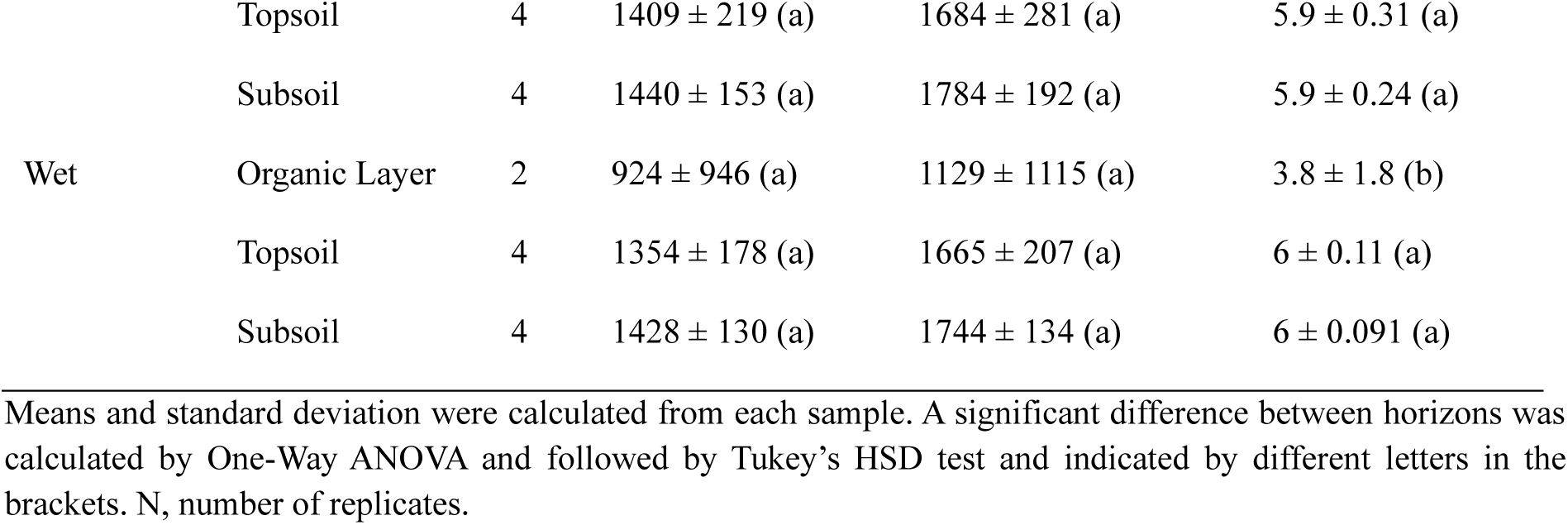
Alpha diversity indices of Bacterial communities.

### Bacterial community composition

The bacterial zOTUs were assigned to 30 different phyla, 230 classes 307 families and 464 genera. The most dominant phyla consisted of Pseudomonadota (24%), Acidobacteriota (16%), Actinobacteriota (15%), Chloroflexota (11%), Verrucomicrobiota (9%), and Gemmatimonadota (7%) (Figure 3). Less abundant phyla, which together accounted for 10% in total were Desulfobacterota, Bacteroidota, Methylomirabilota, Myxococcota, and Planctomycetota. The remaining 19 phyla represented less than 1% of relative abundance. The relative proportions of most bacterial taxa were significantly different among the sites (Figure S3 and S5). Pseudomonadota was dominant in wet and intact site, while the Actinobacteriota was dominant in dry site. At the class level, dry site was dominant by Thermoleophilia (14%) and Acidobacteriae (13%), while wet site was dominant by Gammaproteobacteria (23%) and Gemmatimonadetes (8%). Intact site was dominant by Verrucomicrobiae (15%) and Gammaproteobacteria (14%). The relative proportions of bacterial taxa were also significantly varied across the different horizons of the dry, wet and intact sites (Figure 3 and S4a). The relative proportions of Pseudomonadota, Bacteroidota, Acidobacteriota decreased from the organic layer (or from the topsoil in case of Acidobacteriota) to the subsoil in the dry and wet sites. Similar pattern was also found at class level, like the relative proportions of Gammaproteobacteria, Alphaproteobacteria and Acidobacteriae decreased with the depth in both dry and wet sites. Conversely, the class Thermoleophilia, and Gemmatimonadetes and classes of Chloroflexota increased with increasing depth in these sites. The relative proportions of other classes such as Blastocatellia, Holophagae, Vicinamibacteria and Methylomirabilia were significantly higher in the subsoil compared to other horizons in dry site, and in both topsoil and subsoil compared to the organic layer in the wet sites (Figure 3 and S4b). In case of the intact site, the relative proportions of some bacterial phyla varied significantly between the organic layer and the topsoil and subsoil, but there was less variation in relative proportion between the topsoil and subsoil.

**Figure 3:**
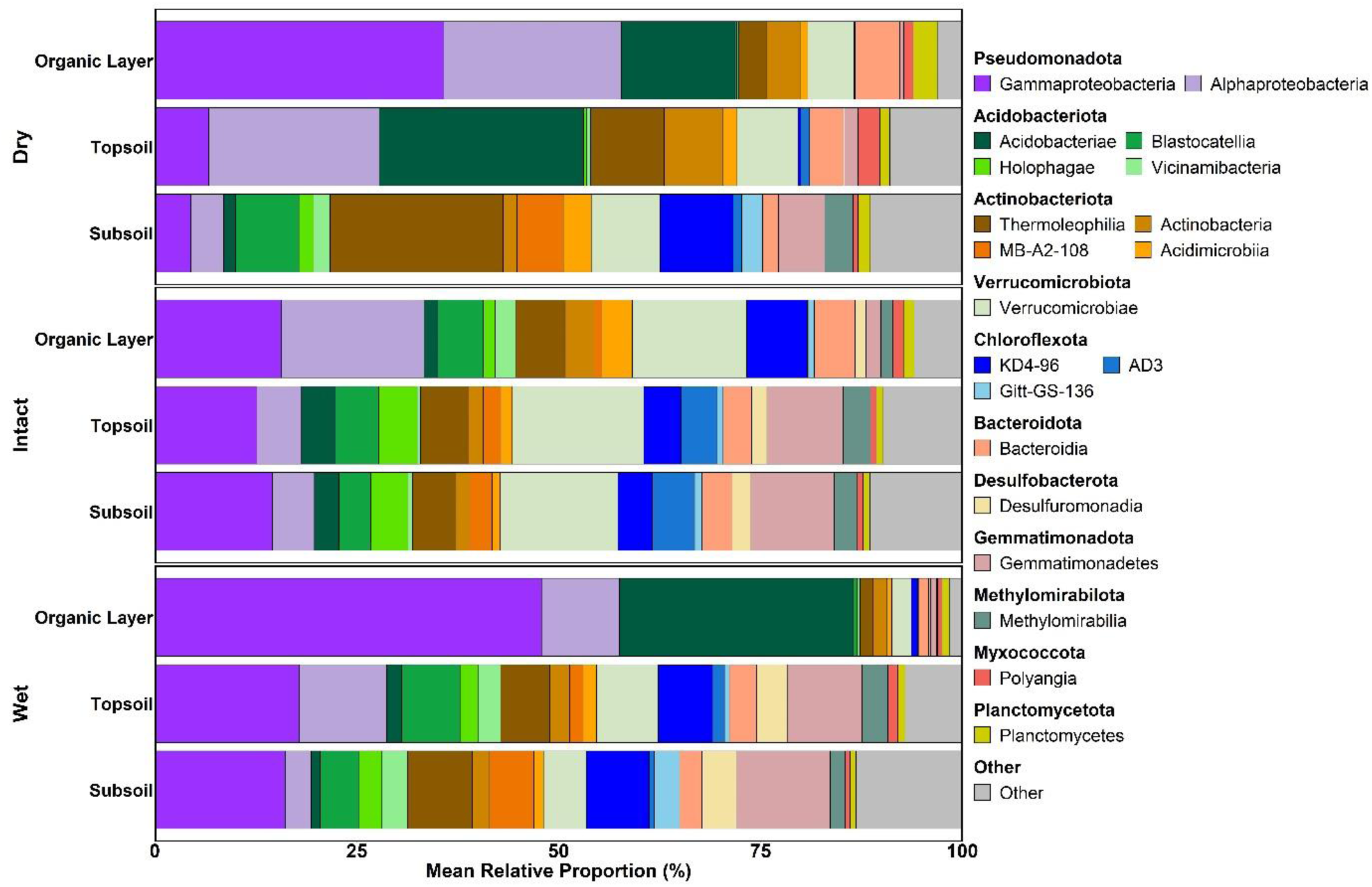
Mean relative proportion of total bacterial community at phylum/class level in in different horizon of dry, wet and intact sites. Taxa representing less than 1% of the total bacterial community are grouped under ‘Other’.

### Bacterial community pattern and relationship with soil parameters

Non-metric multidimensional scaling (NMDS) based on Bray-Curtis dissimilarity was performed to assess the pattern of bacterial communities across studies sites and the three horizons of the dry, wet, and intact sites (Figure 4a and S2). Topsoil and subsoil samples from the wet and intact sites were closely clustered together, while the dry site generally occupied separate place for the topsoil and subsoil. According to the permutation multivariate analysis of variance (PERMANOVA), both sites (F = 4.3, R^2^ = 0.25, p-value = 0.001) and horizon (F= 3.2, R^2^ = 0.2, p-value = 0.001) had strong effect on the beta diversity of bacterial community composition. A redundancy analysis (RDA) based on forward selection was performed to see the relation of soil physicochemicals parameters with the relative proportion of bacterial community (Figure 4b). According to forward seletion, eight parameters were shown signifcant effect on the bacterial community composition. The first two axes of RDA explained 19.8% and 11.4% of the total variation in the community. The bacterial communities in the organic layer were positively correlated to the SOC, C:N, β-glucosidase and phosphatase, but negative correlated to pH. On the other side, bacterial community in the topsoil and subsoil of both wet and intact sites were positively correlated to pH but negatively correlated SOC and C:N, β-glucosidase and phosphatase. N was also selected by forward sletction, however, it was not significantly correlated with bacterial community.

**Figure 4:**
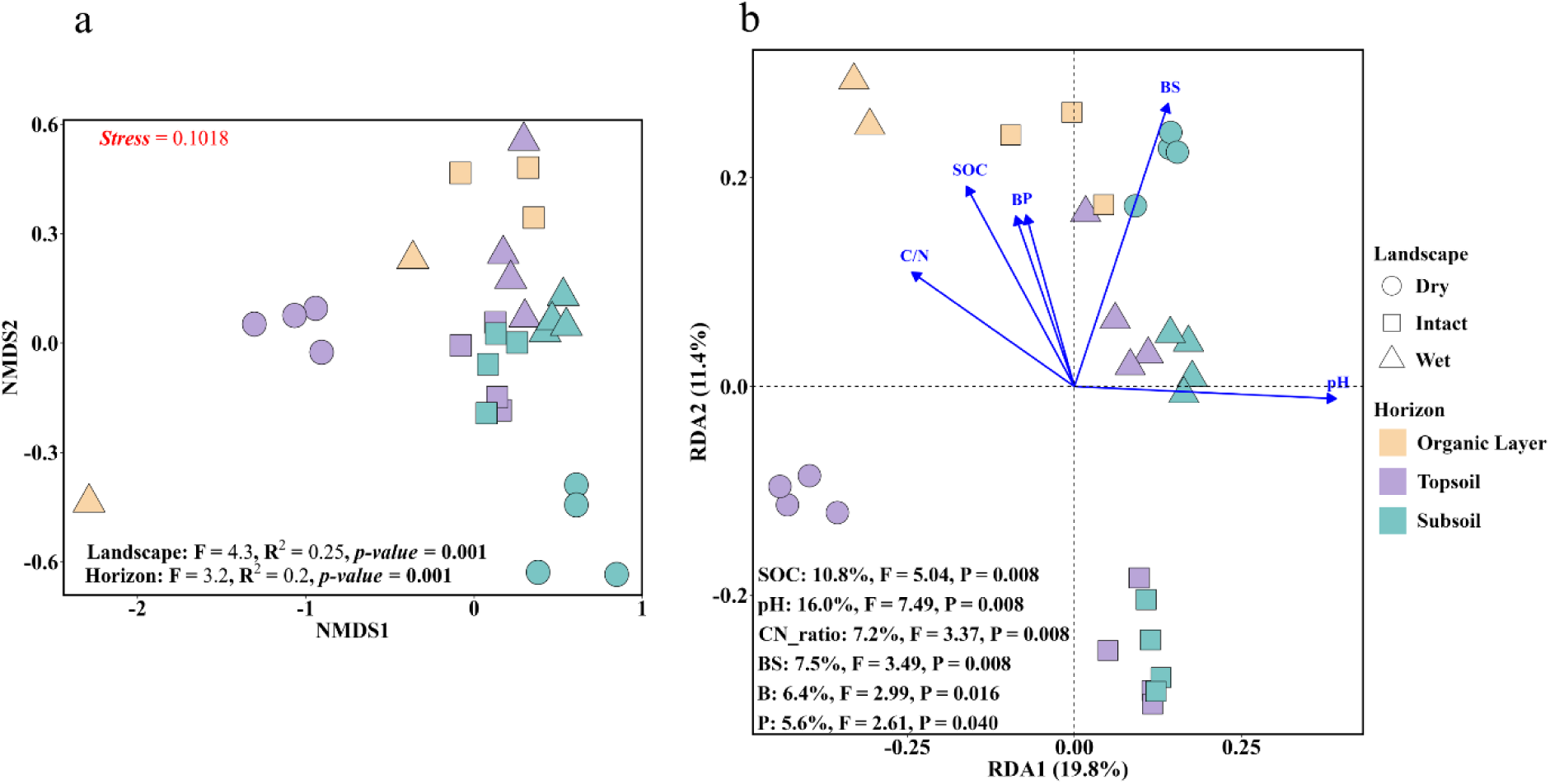
Bacterial community composition pattern. **(a)** Non-metric multidimensional scaling (NMDS) ordination with permutation multivariate analysis of variance (ADONIS: PERMANOVA) of relative proportion of bacterial communities at zOTUs level across the horizon of dry, wet and intact sites based on Bray-Curtis dissimilarities. **(b)** RDA ordination diagram with CCA ANOVA analysis between main soil characteristics (z-score standardized) and changes in the square root-transformed relative proportion of total bacterial communities at zOTUs level. Each point represents a sample, coloured according to horizon and shaped according to sites. The blue arrows represent soil parameters with a significant impact on the structure of bacterial communities. The proportion of variability explained by these parameters is indicated in the bottom-left. SOC: Soil organic carbon, BS: Base saturation, B: β-glucosidase, P: Phosphatase.

According to Spearman correlation, the relative proportion of bacterial taxa was significantly positive or negative effected by the soil physicochemical parameters in the dry and wet degraded permafrost soils compared to intact permafrost soil, where less significant correlations were found (Figure 5). Alphaproteobacteria, Gammaproteobacteria, and Acidobacteriota were positively correlated with SOC, N, C:N, moisture, EPS-protein and EPS-sugar, but negatively correlated with pH and BS in the dry site and with only pH in the wet site (Figure 5a). In contrast, Methylomirabilota, Chloroflexota, Gemmatimonadota, Actinobacteriota, and Nitrospirota were negatively correlated with all parameters except pH and BS, where they were positively correlated with pH and BS in the dry site and with only pH in the wet site. Bacteroidota was positively correlated with all parameters except the pH and BS in dry site, while negatively correlated with most parameters except the pH in wet site. In the intact site, the effect of soil parameters on bacterial taxa were not prominent compared to the dry and wet sites but some correlations were found (Figure 5a). For example, Planctomycetota was significantly positively correlated with SOC, N, and moisture in intact site, but no significant correlations were observed in the dry and wet site. Actinobacteriota was positively correlated with moisture in intact site, whereas the correlation with moisture was strongly negative in dry and wet sites. Similarly, Methylomirabilota was negatively correlated with pH and BS in intact site, while it was positively correlated with pH and base saturation in dry and wet sites.

**Figure 5:**
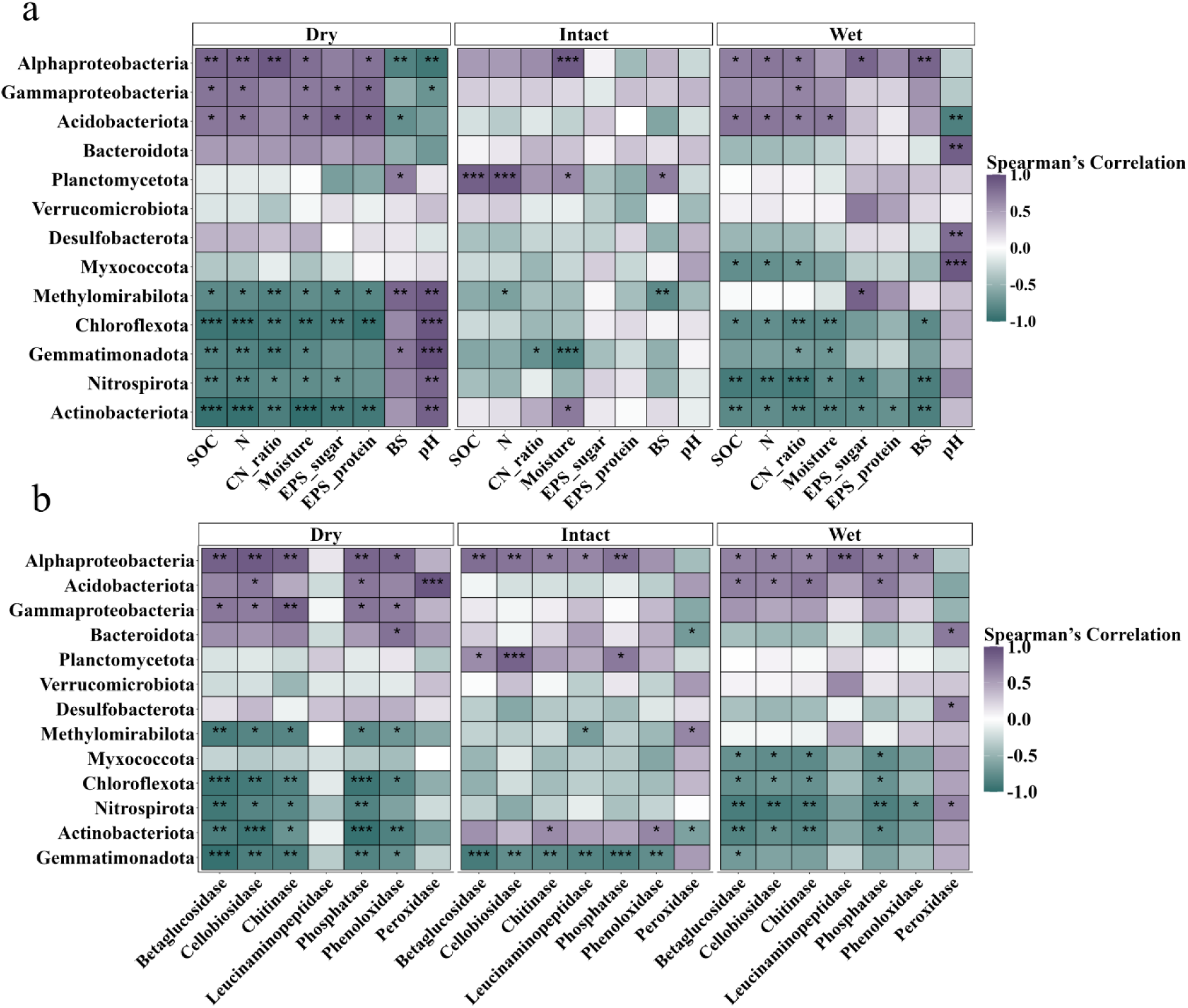
Heatmap visualizing the Spearman’s correlation values between the relative proportion of bacterial communities at phylum/class level and (a) soil physicochemical parameters and (b) soil enzymatic activities across the dry, wet and intact sites. The color intensity of each cell indicates the strength and direction of the correlation, ranging from dark green (negative correlation) to dark purple (positive correlation). White cells represent correlations close to zero. The significance of each correlation is indicated within the cells by *P_Value* 0.001, “***”, *P_Value* 0.01 “**”, *P_Value* 0.05 “*”.

A strong correlation also existed between the relative proportions of bacterial taxa and the abundance of extracellular enzymes (Figure 5b). Alphaproteobacteria, Gammaproteobacteria, and Acidobacteriota were positively correlated with β-glucosidase, cellobiosidase, chitinase, phosphatase, PhOx and PerOx in the dry site. Similar positive correlation was also found in the wet site except for the PerOx, where Alphaproteobacteria, Gammaproteobacteria and Acidobacteriota were negatively correlated with PerOx. In the intact site, Alphaproteobacteria was positively correlated with most of the enzymes, while the Gammaproteobacteria and Acidobacteriota were not significantly correlated with enzymes. Chloroflexota, Nitrospirota, Gemmatimonadota, and Actinobacteriota were negatively correlated with β-glucosidase, cellobiosidase, chitinase, phosphatase, PhOx and PerOx in the dry site as well as in the wet site with the only exception of their positively correlation with PerOx. In the intact site, the Gemmatimonadota was negatively correlated with all enzymes except for the PerOx, whereas Actinobacteriota showed the opposite pattern in their correlations. Planctomycetota was significantly positively correlated with cellobiosidase, chitinase and phosphatase in the intact site, while no significant correlation was found in dry and intact site.

## Discussion

### Bacterial diversity patterns under hydrological conditions of degraded permafrost soil

Previous studies underscore the role of physicochemical processes in varied conditions of degraded permafrost soils (Lee et al., 2012; Liebmann et al., 2024; Schädel et al., 2016; Song et al., 2020; Jorgenson et al., 2013; Treat et al., 2016) and microbial communities mostly in waterlogged soils (Coolen & Orsi, 2015; Fahle et al., 2014; Wu et al., 2018). In this study, we examined total bacterial community abundance and composition, along with their relationship to soil physicochemical factors, across two contrasting hydrological landscapes of degraded permafrost soils; dry oxic soil and wet anoxic soil, comparing these to non-degraded intact permafrost soil. Our results showed significantly higher bacterial alpha diversity (Observed and Chao1) in wet and intact sites than in dry site, suggesting similar bacterial richness between wet and intact sites but lower richness in dry site (Figure 2). Similarly, a higher proportion of shared zOTUs was observed between wet and intact sites then the dry site (Figure 1). This finding supports our first hypothesis that the bacterial community structure in wet degraded permafrost soil is more closely resembling that of intact permafrost soil. This is likely due to the shared physicochemical traits such as water saturation and higher SOC content between the wet and intact sites (Liebmann et al., 2024). In contrast, dry site exhibits increased aeration and oxic conditions that enhance microbial activities and processes related to OM decomposition and SOC loss (Kwon et al., 2019; Natali et al., 2015; Jorgenson et al., 2013). Therefore, these conditions in dry site along with pronounced leaching structures visible in the soil profiles, less prevalent in wet and intact sites (Liebmann et al., 2024), can contribute to the shift in bacterial community structure.

The relatively high OC content and nutrient inputs from the ground surface vegetation in the topsoil support a variety of bacteria adapted to drier, oxidizing environments and leading to enhanced bacterial richness and diversity (Ren et al., 2018). Studies have consistently shown that bacterial alpha diversity is higher in the topsoil compared to the subsoil horizon of permafrost soils (Chen et al., 2017; Ren et al., 2022). This trend aligns with our findings only for the dry site, where the alpha diversity found higher in the mineral topsoil then the mineral subsoil (Table 3). The opposite trend was found in the wet site, where alpha diversity was higher in the subsoil than in the topsoil. The higher alpha diversity in the subsoil of the wet site may be more associated with decomposition of previously stored OM and nutrient cycling, leading to lower C:N ratio due to rapid N cycling and OM turnover. The study by Kuhry et al. (2020) supports these findings, indicating that soils with lower C:N ratios tend to have higher microbial activity and more efficient nutrient cycling, which aligns with the observed higher alpha diversity in the subsoil of the wet site.

### Bacterial taxonomic richness in degraded permafrost soil

In our second hypothesis, we predicted that bacterial taxonomic richness in the wet site would exhibit reduced diversity, similar to that observed in intact permafrost soils. This assumption was based on the idea that excessive moisture could restrict microbial adaptability and limit diversity due to hypoxic conditions, reducing the number of taxa that are able to thrive. However, contrary to our hypothesis, we observed no significance difference in overall taxonomic richness at phylum/class level between wet and dry sites, suggesting that moisture may not be a primary factor driving bacterial taxonomic structure in these landscapes (Figure 3 and S5). This finding is further supported by the results of redundancy analysis with forward selection, which did not identify moisture as a significant explanatory variable for bacterial community composition (Figure 4b). Instead, our results showed only a significant variation in the relative proportions of bacterial taxa between these landscapes (Figure S3). Bacterial composition and abundance are also greatly affected by the depth due to variations in pH, moisture, C and N contents (Baker et al., 2023; X. Wu et al., 2017). The abundance of most bacterial taxa was significantly different across the horizons of studied sites with this pattern being more pronounced in the dry and wet degraded permafrost soil than the intact permafrost soil (Figure 3 and S4). This significant variation may result from the larger distance between horizons mainly mineral topsoil and subsoil (about 45 cm) in both dry in wet sites compared to the intact site where the distance was about 10 cm (Liebmann et al., 2024). We found that classes Acidobacteriae and Thermoleophilia were significantly dominant in the dry site, while Gemaprotobacteria was dominant in the wet site. The high abundance of Acidobacteriae is primarily associated with the low pH (Kobabe et al., 2004; Männistö et al., 2007) and limited OC content as they are considered oligotrophic (Eichorst et al., 2011; B. Yan et al., 2021). Studies suggest that although Acidobacteriae are generally considered oligotrophic, certain members of this phylum also exhibit high relative abundances even in environments with elevated OC content (Bárta et al., 2017; Jones et al., 2009). Similar in our study, the relative proportion of Acidobacteriae was higher in the OC-rich organic layer and topsoil compared to the subsoil in the dry site where OC content was more limited. This observation indicates that factors beyond just C availability such as N or specific OM types may play a significant role in driving the metabolism and ecological success of the member of this class (Bárta et al., 2017). However, the influence of pH on the abundance of Acidobacteriae appears particularly significant, as they were less abundant in the topsoil (more alkaline) of the wet site than the topsoil (more acidic) of the dry site. Thermoleophilia, on the other hand, were significantly abundant in the subsoil than the topsoil and organic layer of both dry and wet sites (Figure 3 and S4). They are associated with oligotrophic environments where they excel as slow-growing, adapted to low-nutrient conditions and complex C substrates (Ricketts et al., 2020; Y. Yang et al., 2023). Therefore, their abundance in the subsoil, particularly at the dry site, suggests that they can outcompete the copiotrophic groups like Alphaproteobacteria and Gammaproteobacteria, due to their ability to decompose more stable OM, such as aliphatic and aromatic compounds (Shi et al., 2020; Sun et al., 2021; Y. Yang et al., 2023), which are less accessible to fast-growing copiotrophs. Some taxa such as anaerobic methanotrophs (Methylomirabilia) and sulfur reducers (Desulfuromonadia) were uniquely abundant across all horizons of the wet and intact sites, whereas they were undetected or restricted to the subsoil in the dry site. Members of Methylomirabilia are known for their ability to oxidize CH₄ even in oxygen-free environments by producing oxygen internally through nitrite reduction (Ettwig et al., 2010). Their presence in the deep horizons of wet anoxic condition suggest that they may be contributors to the OM decomposition, by coupling CH₄ oxidation with denitrification, thus playing a critical role in C and N cycling in these (temporarily) anoxic environments (Yao et al., 2024). While members of Methylomirabilia are key players in CH₄ oxidation and OM decomposition in deep anoxic horizons, the active layer of the intact site was significantly dominated by methanotrophs Verrucomicrobiae from the Verrucomicrobiota phylum may act as potential contributors to CH₄ emissions during warmer seasons due to their capacity for CH₄ oxidation (Op den Camp et al., 2009). However, some of the strains from Verrucomicrobiota have been identified as having N-fixing capabilities, harboring nifH genes essential for N fixation (Penton et al., 2016; Wertz et al., 2012). This is particularly significant given that N is often a limiting nutrient in permafrost soils and may help alleviate N limitation mainly in the active layer of intact permafrost soil.

### Influence of soil physiochemical factors on bacterial community abundance and structure

Bacterial community structures (beta-diversity) strongly differed between the studied sites and also differed along the different horizons (Figure 4a and S2). These differences were strongly driven by the differences in soil physicochemical parameters, which might largely be attributed to differences in hydrological conditions (Figure 4b). Soil pH plays a crucial role in shaping bacterial community structure across various ecosystems, including Arctic permafrost regions (Chu et al., 2010; Fierer & Jackson, 2006; Ren et al., 2018). The lower pH in the topsoil of the dry site (4.5) compared to the wet and intact sites (around 5.9) may be attributed to the accumulation of acidic compounds from OM decomposition (Li et al., 2008; F. Yan et al., 1996), creating a favourable environment for acidophiles. In contrast, the higher pH in the topsoil of the wet and intact sites could result from the slower OM turnover under water-saturated conditions, which limits organic acid production and maintains a relatively stable or higher pH (Yan et al., 1996), thereby supporting the presence of alkaliphilic bacterial taxa. These findings are consistent with the observed strong negative correlation between the relative proportion of bacterial communities and pH in the dry site topsoil, whereas bacterial communities in the wet and intact topsoil showed a positive correlation with pH (Figure 4b). In addition to pH, SOC, C:N ratio, and the activities of key extracellular enzymes such as β-glucosidase and phosphatase significantly influenced bacterial composition and abundance across the dry, wet, and intact sites. Higher SOC concentrations, particularly in surface horizons, provided labile C substrates that likely favoured copiotrophic taxa, whereas lower SOC in subsoil appeared to support oligotrophic taxa adapted to nutrient-poor conditions (Li et al., 2021). Similarly, variations in the C:N ratio influenced bacterial taxa by altering N availability, favouring N-fixing and N-utilizing microbes in horizons with lower ratios (Kuhry et al., 2020). Notably, the relative proportion of bacterial taxa in the organic layer of the wet and intact sites which are mainly abundant with copiotrophs showed a positive correlation with SOC and the C:N ratio, while those in the subsoil mostly oligotrophs exhibited a negative correlation. From the correlation analysis, we found that the relative proportion of dominant classes of copiotrophic Pseudomonadota, and Acidobacteriota in the organic layer and topsoil of both dry and wet degraded permafrost soil were strongly positively correlated with SOC and C:N ratio, while the relative proportion of oligotrophic taxa found more abundant in the subsoil such as Actinobacteriota, Gemmatimonadota and Methylomirabilota displayed a negative correlation (Figure 5a). This strong correlation patterns were more prominent in the dry and wet sites compared to the intact site as we suggested in our third hypothesis. This could be attributed to the increased soil temperatures in the degraded landscapes, as opposed to the intact permafrost soil (Y. Song et al., 2022). Another reason could be the accumulation of labile C sources in the upper horizon, derived from well-established vegetation, or the release of OC at deeper horizons previously sequestered in permafrost soil, may have stimulated bacterial activity (Coolen et al., 2011; Pautler et al., 2010; Schuur et al., 2008). Furthermore, the degradation of permafrost alters soil structure, improving conditions for bacterial colonization and activity (Z. Yang et al., 2010). These structural changes, such as enhanced soil porosity and aeration, likely promote bacterial processes, further reinforcing the observed correlations in both dry and wet degradation landscapes.

Furthermore, permafrost degradation has a significant impact on soil enzyme activities due to changes in temperature, moisture, OM availability and soil depth (Coolen & Orsi, 2015; Schuur et al., 2015; Y. Song et al., 2023). Our findings indicated higher extracellular enzyme activities in both dry and wet degraded permafrost soil than the intact permafrost soil site (Table S2). This increase may be due to the higher levels of SOC in the upper horizons, which are generally more labile and easily mineralizable (Piotrowska-Długosz et al., 2022) as well as the exposure of previously frozen OM (Coolen & Orsi, 2015). Therefore, our findings of increased enzymatic activities and their correlation with relative proportion of bacterial taxa in both dry and wet degraded permafrost soils may align closely to the SOC loss and OM decomposition. For instance, the positive correlation of Pseudomonadota and Acidobacteriota with hydrolytic enzymes (β-glucosidase, cellobiosidase, and chitinase) suggests that the members of these taxa may play a crucial role in decomposing OC in both degradation landscapes (Figure 5b). This observation is consistent with previous studies in thawed permafrost regions, where enhanced microbial activity has been linked to significant C release, particularly through CO₂ and CH₄ emissions (DeConto et al., 2012; Koven et al., 2011; Schuur et al., 2008). These results suggest that accelerated bacterial decomposition, driven by high enzymatic activities can be a key factor in the loss of SOC stock from degraded permafrost soil either dry oxic or wet anoxic conditions.

However, a recent study reported higher SOC stocks and a greater abundance of macroaggregates in the topsoil of both dry and wet degraded permafrost soils compared to intact permafrost soils (Liebmann et al., 2024). These findings can also be linked to the enhanced production of EPS, primarily by EPS-producing bacteria, which promote soil aggregation by binding soil particles into larger aggregates (Costa et al., 2018; Mueller et al., 2017). Our results align with this observation, as we found higher levels of EPS (both polysaccharide content, measured as glucose equivalent, and EPS-protein) in the topsoil of the dry and wet sites compared to the intact site (Table 2). These elevated EPS levels likely contribute to the formation of macroaggregates and are consistent with the higher SOC stocks observed in degradation sites, indicating the important role of EPS in improving soil structure and stabilizing OC. Furthermore, we observed a positive correlation between EPS content and the relative proportions of Alphaproteobacteria, Betaproteobacteria, and Acidobacteriota (Figure 5b). Specific genera within these groups, such as *Rhizobium* and *Sphingomonas* (Alphaproteobacteria) (Alami et al., 2000; Azeredo & Oliveira, 2000), and *Burkholderia* (Betaproteobacteria) (Harahap et al., 2018), are well-documented producers of EPS that facilitate soil aggregation by binding soil particles and creating stable macroaggregates. This suggests that the presence of these taxa in certain proportion in both degradation sites could play an active role in EPS production and are key contributors to the observed soil aggregation and over all stability of SOC by occlusion of OM within aggregates.

## Conclusion

Permafrost degradation triggers significant shifts in bacterial communities, driven by increased soil temperatures, moisture changes, and nutrient availability. The findings of this study emphasize that hydrological and physicochemical variations in degraded permafrost environments profoundly influence bacterial community structure and their abundance. The bacterial alpha diversity in the wet degraded permafrost closely mirrors that of intact permafrost, likely due to similar hydrological and soil characteristics such as water saturation and high SOC content. Conversely, the dry degradation landscape, with its more aerated and oxic conditions, supports distinct bacterial communities from the other two studied sites, demonstrating the significant influence of soil physicochemical conditions on bacterial diversity and community composition across the degraded permafrost landscapes. If it comes to the higher taxonomic rank, we observed no significant difference in bacterial structure, but their relative proportion was significantly differed between dry and wet sites, and across the horizons. They were largely influenced by soil physicochemical parameters like pH, SOC, and the C:N ratio, however moisture level was not the significant factor to influence the bacterial taxa abundance . SOC promoted copiotrophic taxa in upper horizons, while nutrient-poor or complex OC substrate subsoils supported the oligotrophs. The presence of aliphatic degraders (Thermoleophilia), acidophilic and cellulolytic (Acidobacteriae) together with the copiotrophs and other oligotrophs may be the potential candidates for the OM decomposition and SOC loss. While in the wet site anaerobic methylotrophs such as members of Methylomirabilota and Verrucomicrobiota could be main contributors in the OM decomposition and SOC loss in the form of CH₄. Additionally, the higher EPS levels in both degraded sites, produced by bacterial taxa, as we found a strong positive correlation with Alpha and Beta-proteobacteria, and Acidobacteriota likely one of the main sources that facilitate soil aggregation and stabilizing SOC through the occlusion of OM within aggregates. This dual role of bacterial communities in both accelerating SOC loss and improving soil structure demonstrates their adaptive strategies in different permafrost degradation landscapes.

## Data availability

Raw fastq 16S rRNA gene sequences of the whole microbiome were uploaded to European nucleotide archive (ENA) under the study n. PRJEB79237.

## Author contribution

CV, JB, PL, TU and GG completed fieldwork and collected samples from Fairbanks, Alaska, USA. MW, HW and MV performed molecular analysis and JB and MN analzsed the sequencing data. CV extracted EPS from the soil samples and determined into EPS polysaccharide and protein. PL determined all the soil physicochemical parameters. MW and JB wrote the manuscript, and CV, GG, HW, TU, and OS contributed to and have approved the final manuscript.

