## Supplementary material for "Different hydrological conditions after permafrost thaw result in distinct microbial community compositions": s

Table S1: Soil physiochemical properties

| **Locality** | **Count** | **Moisture [%]** | **pH** | **BS [%]** | **SOC [mg g$^{-1}$]** | **N [mg g$^{-1}$]** | **C:N** | **EPS_sugar [µg g$^{-1}$]** | **EPS_protein [µg g$^{-1}$]** |
|---|---|---|---|---|---|---|---|---|---|
| Dry | 9 | 33.2±19.5(a) | 5.3±1(a) | 48.2±20.8(a) | 59±128.8(a) | 2.6±5(a) | 15.6±6.6(a) | 58.4±29.8(a) | 30.2±18.9(a) |
| Intact | 11 | 52.4±26.9(a) | 5.9±0.5(a) | 57.1±20.8(a) | 116.9±166.1(a) | 4.2±5(a) | 19.7±9.9(a) | 69±28.9(a) | 34.5±28.9(a) |
| Wet | 10 | 43.3±19.1(a) | 5.6±0.8(a) | 61.2±10.2(a) | 85.9±142.3(a) | 3.2±4.4(a) | 17.4±9.4(a) | 134.8±128.3(a) | 33.7±27.4(a) |

Averages and standard deviation were shown. The significant difference between horizons was calculated by One-Way ANOVA and followed by Tukey's HSD test. Different letters in the brackets indicate a significant difference. Count: number of replicates; na, no data available. BS: Base saturation, SOC: Soil organic carbon, N: Nitrogen

Table S2: Soil enzyme activity per landscape

| **Landscape** | **Count** | **Betaglucosidase [nmol g$^{-1}$ dw h$^{-1}$]** | **Cellobiosidase [nmol g$^{-1}$ dw h$^{-1}$]** | **Chitinase [nmol g$^{-1}$ dw h$^{-1}$]** | **Phosphatase [nmol g$^{-1}$ dw h$^{-1}$]** | **Leucinaminopeptidase [nmol g$^{-1}$ dw h$^{-1}$]** | **Phenoloxidase [µmol h$^{-1}$ g$^{-1}$dw]** | **Peroxidase [µmol h$^{-1}$ g$^{-1}$dw]** |
|---|---|---|---|---|---|---|---|---|
| Dry | 9 | 280.9±580.5 (a) | 88.1±213.6 (a) | 175.4±435.8 (a) | 482.1±1132.8 (a) | 17.6±9.8 (a) | 152.6±180.5 (a) | 735.7±76.4 (b) |
| Intact | 8 | 58.7±27.3 (a) | 12.7±3.8 (a) | 26.2±11.7 (a) | 70.8±27.1 (a) | 21±6.1 (a) | 87.2±26.2 (a) | 1083.2±142.1 (a) |
| Wet | 10 | 231.4±374.7 (a) | 22.3±30.1 (a) | 72.2±80.6 (a) | 326.2±436.1 (a) | 50.7±51.1 (a) | 151±75.3 (a) | 1042.2±403.3 (ab) |

Averages and standard deviation were shown. The significant difference between horizons was calculated by One-Way ANOVA and followed by Tukey's HSD test. Different letters in the brackets indicate a significant difference. Count: number of replicates.

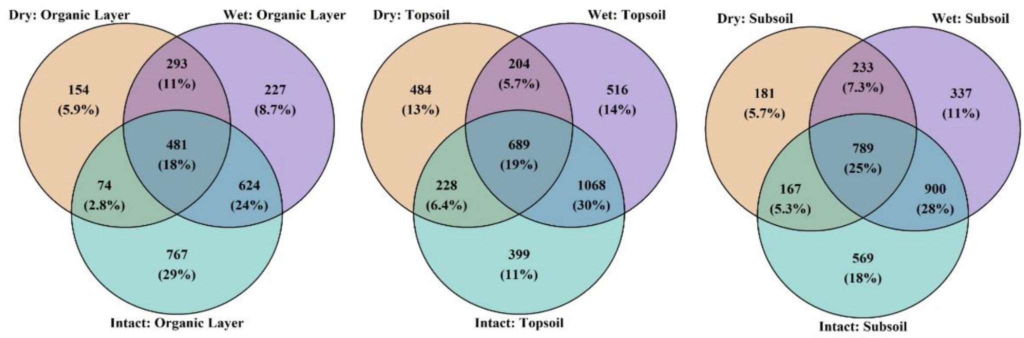


**Figure S1:** Venn diagram illustrating the overlap of zOTUs across the horizons of the dry, wet, and intact landscapes. The diagram displays both the raw counts and the percentages of zOTUs unique to each condition and shared among them, with the sizes of the circles representing the total number of zOTUs in each condition. The intersections highlight the shared zOTUs between the conditions, with the central area representing the common zOTUs.

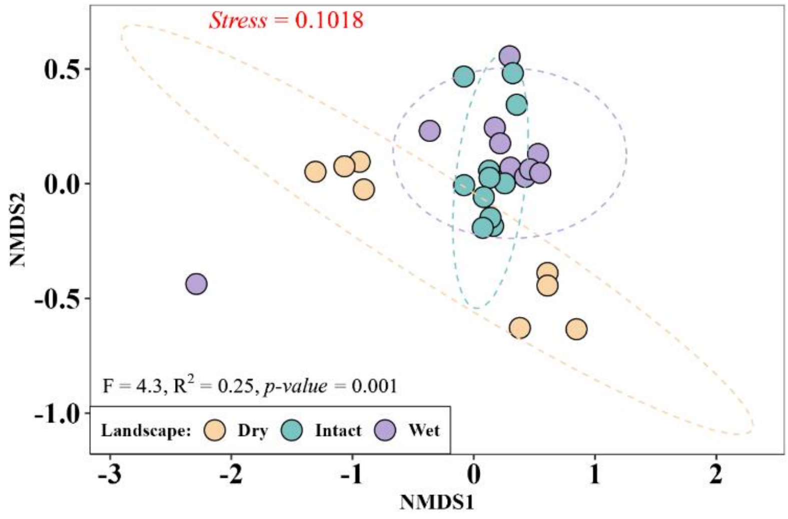


**Figure S2: NMDS plot** based on **Bray-Curtis distance**, illustrating the ordination of **bacterial communities** across different **Sites**. Each point represents a sample, colored according to its locality, and (B) ellipses indicate the 95% confidence intervals around the centroid of each locality group. The Analysis of Similarities **(ADONIS)** was conducted to assess the differences in bacterial community composition among the localities. The global **ADONIS** result indicates a significant difference in community composition (ADONIS: F Model= 4.3, R2 = 0.25, *p-value* = 0.001). Pairwise ADONIS comparisons were also performed between localities to further investigate these differences. Outliers were identified as samples with distances from the centroid greater than the 95th percentile (threshold = 0.95) and were subsequently removed from the analysis.

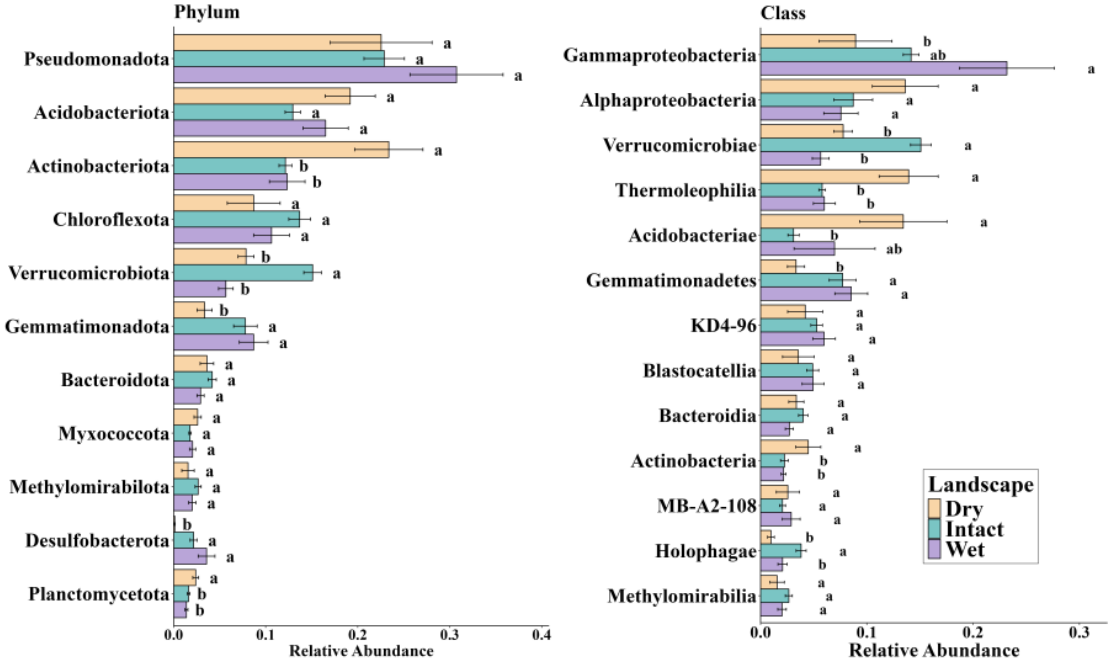


**Figure S3:** Relative abundance of bacterial phyla across the dry, wet and intact landscape. The analysis was conducted using one-way ANOVA to assess differences in bacterial taxa abundance among the landscapes. The significance of differences is indicated by letters on the bars, with lowercase letters denoting significant differences ($p < 0.05$).

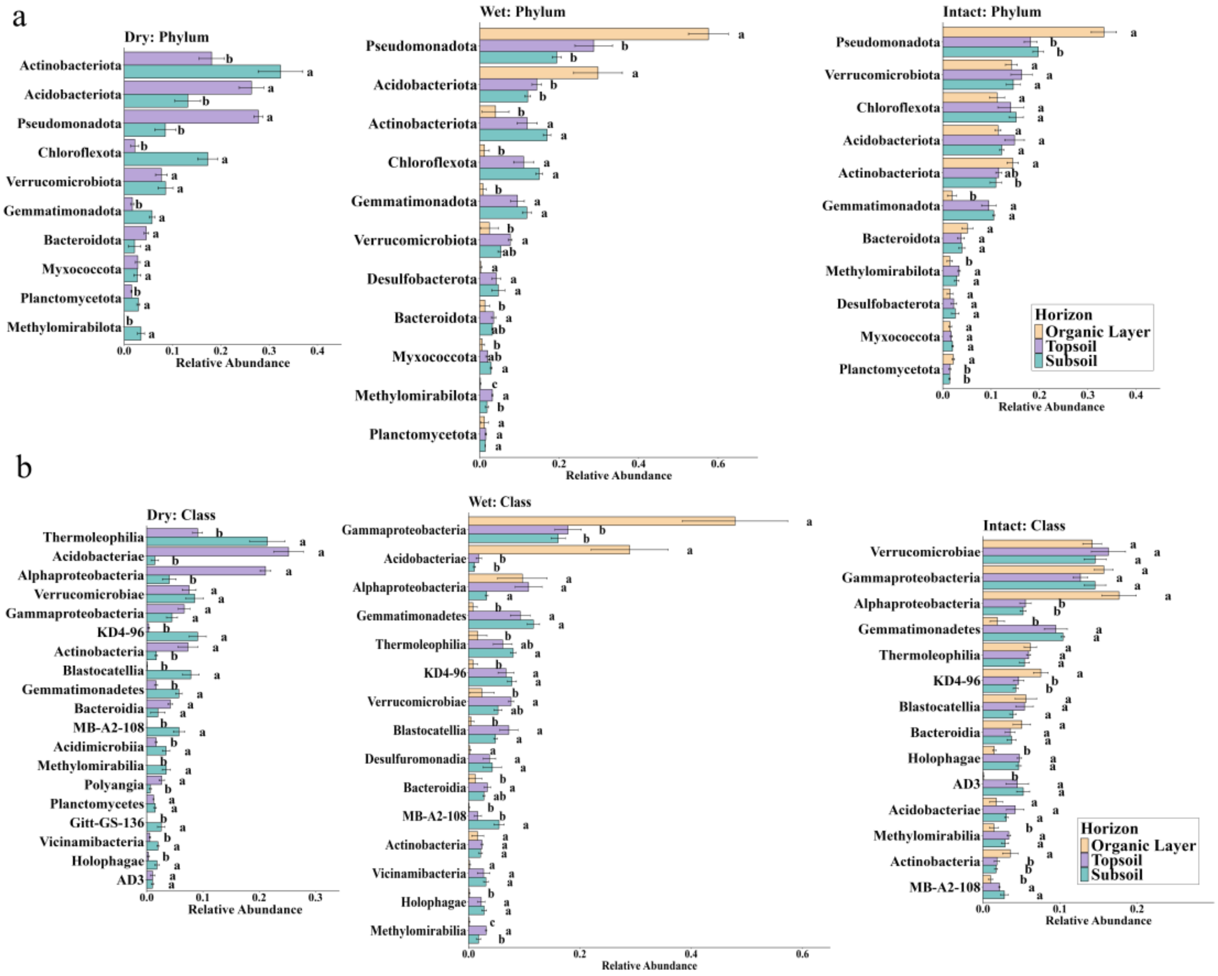


**Figure S4:** Relative abundance of bacterial (a) phyla and (b) class across different horizons of dry, wet and intact landscape. One-way ANOVA was performed to assess differences in bacterial taxa abundance among the horizons. The significance of differences between horizons is indicated by letters on the bars, with lowercase letters denoting significant differences ($p < 0.05$).

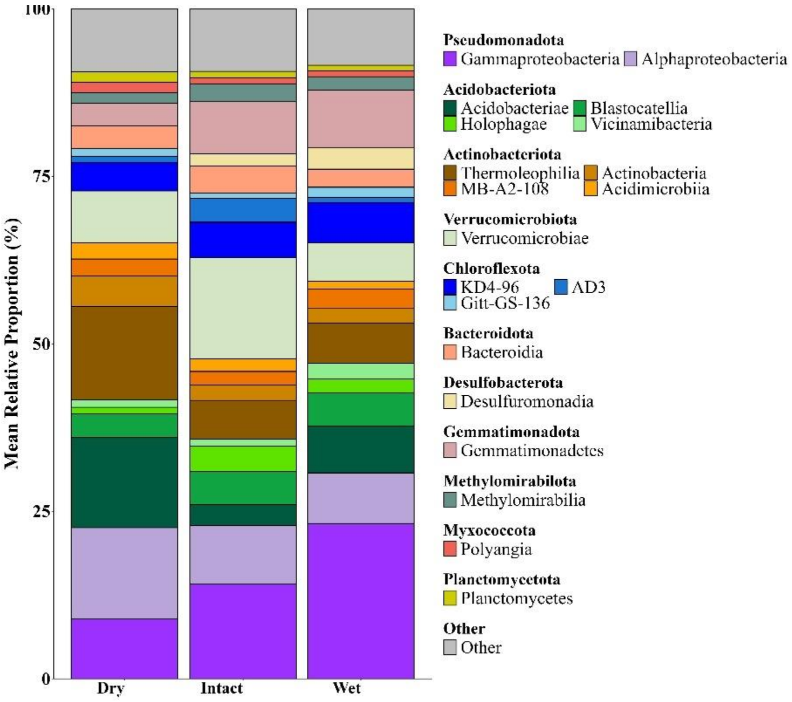

**Figure S5:** Mean relative proportion of total bacterial community at phylum/class level in dry, wet and intact landscape. Phyla representing less than 1% of the total bacterial community are grouped under 'Other'.
